# PReMS: Parallel Regularised Regression Model Search for sparse bio-signature discovery

**DOI:** 10.1101/355479

**Authors:** Clive J. Hoggart

## Abstract

There is increasing interest in developing point of care tests to diagnose disease and predict prognosis based upon biomarker signatures of RNA or protein expression levels. Technology to measure the required biomarkers accurately and in a time-frame useful to health care professionals will be easier to develop by minimising the number of biomarkers measured. In this paper we describe the Parallel Regularised Regression Model Search (PReMS) method which is designed to estimate parsimonious prediction models. Given a set of potential biomarkers PReMS searches over many logistic regression models constructed from optimal subsets of the biomarkers, iteratively increasing the model size. Zero centred Gaussian prior distributions are assigned to all regression coefficients to induce shrinkage. The method estimates the optimal shrinkage parameter, optimal model for each model size and the optimal model size. We apply PReMS to six freely available data sets and compare its performance with the LASSO and SCAD algorithms in terms of the number of covariates in the model, model accuracy, as measured by the area under the receiver operator curve (AUC) and root predicted mean square error, and model calibration. We show that PReMS typically selects models with fewer biomarkers than both the LASSO and SCAD algorithms but has comparable predictive accuracy.

**Availability:** (PReMS) is freely available as an R package https://github.com/clivehoggart/PReMS

## 1. Introduction

For many diseases clinical presentation features alone cannot reliably be used for diagnosis, this has motivated the development of proteomic and transcriptomic signatures, for example, to diagnose tuberculosis (TB) (Kaforou *et al.* (2013); Anderson *et al.* (2014); Chegou *et al.* (2016)), Kawasaki disease (Jaggi *et al.* (2018)) and to distinguish bacterial from viral infections (Herberg *et al.* (2016)). All such biosignatures are derived from comparing cases and controls with measures of potential biomarkers; ranging from tens of proteins from a multiplex cytokine platform to tens of thousands from whole genome transcriptome analysis. Logistic regression with biomarker selection via simple forward selection or penalised regression in the form of the LASSO (least absolute shrinkage and selection operator) (Tibshirani (1996)) or elastic net (Zou and Hastie (2005)) have been frequently used for biomarker signature estimation (Kaforou *et al.* (2013); Anderson *et al.* (2014); Herberg *et al.* (2016)). Logistic regression is attractive as its implementation simply requires a weighted sum of the biomarkers selected and as such it is easy to interpret, and the LASSO and elastic net both implement biomarker selection and shrinkage of the regression coefficients resulting in more stable model estimation.

A major challenge in using biomarker signatures as diagnostic tools is their translation to clinical tests suitable for use in hospital laboratories or at the bedside. The fewer biomarkers used by a signature the easier it will be for it to be turned into a point of care test (Cattamanchi *et al.* (2013); Gjen *et al.* (2017)). However, both the LASSO and elastic net tend to select large models by allowing some noise predictors to enter the model (Buhlmann and van de Geer (2011)) suggesting similar predictive accuracy could be attained with fewer predictors. This paper is motivated by the task of estimating prognostic and diagnostic biosignature models using a minimal set of markers.

We present the Parallel Regularised Regression Model Search (PReMS) algorithm, implemented in R, for selecting an optimal set of biomarkers, from a set of many potential biomarkers, for logistic regression biosignatires. We demonstrate that the method has similar predictive accuracy as the LASSO whist utilising fewer biomarkers. The method is conceptually straightforward: search as many models as possible and choose the best one; throughout we use model to mean a subset of selected biomarkers with their respective logistic regression coefficients. The implementation utilises the multiple processors available on desktop computers using the R package parallel (Eddelbuettel (2018)) making the search of very large model spaces computationally feasible. The problem can then be split in two: estimating the regression coefficients of each model and 2) criteria for selecting the best model. Gaussian shrinkage priors are applied to the regression coefficients for model robustness. To aid computational efficiency the LASSO, as implemented in the R package glmnet (Friedman *et al.* (2010)), is used to derive an estimate for the Guassian shrinkage parameter and the Watanabe-Akaike Information Criteria (WAIC) (Watanabe (2010)) is used to select the optimal model of a given size.

We apply the method to freely available biological data sets and compare results to those obtained with the LASSO with two alternative penalty parameters and the smoothly clipped absolute deviation (SCAD) (Fan and Li (2001)) algorithm. The SCAD algorithm was included in the comparison as it is known to select relatively sparse models. We do not compare with the elastic net as it is known to select larger models than the LASSO and our focus is on small models.

## 2. Materials and Methods

Regression coefficients can be naively estimated by their maximum likelihood estimates (MLEs). However, in situations in which there are more covariates than observations (*p > n*) the MLE is not identifiable and otherwise results in over-fitting and sub-optimal predictive accuracy (Pavlou *et al.* (2016)). This can be simply remedied by ‘shrinking’ the MLEs towards zero. From a Bayesian perspective shrinkage is achieved by assigning the regression coefficients zero centred symmetric prior distributions, the greater the prior precision (=1/variance) the greater the shrinkage. All model selection methods considered employ shrinkage of logistic regression coefficients. Logistic regression is used as we are interested in prediction of a binary outcome, ie disease status, however, the method is equally applicable to other generalised linear model. The log-posterior, or equivalently the penalised log-likelihood, of logistic regression shrinkage models is

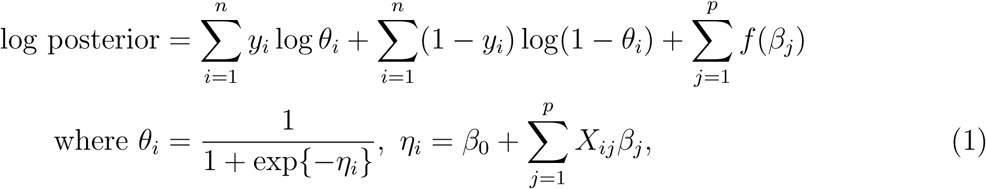

*y_i_* is the outcome of the *i*th individual (*i* = 1*,…, n*) taking values 0 or 1, ***β*** = (*β*_1_*,…, β_p_*) are the regression coefficients, *β*_0_ is the intercept, *X_ij_* is covariate *j* for individual *i*, *θ_i_* is the predicted probability for individual *i* and *f* is the log-prior or minus penalty function. The solution for any given penalty function is given by maximising the log-posterior with respect to ***β***.

### 2.1. LASSO

From a Bayesian perspective the LASSO assigns regression coefficients double exponential prior distributions. With shrinkage parameter *λ* the log-prior can be expressed as

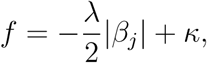

for all coefficients *β_j_*, *κ* is a constant. The greater *λ* the greater the shrinkage. We use the R package glmnet (Friedman *et al.* (2010)) to fit the LASSO and consider two solutions for the penalty parameter provided by the package: one which minimises the leave-one-out cross validated deviance (denoted by *λ_opt_*) and the other derived from the penalty which gives a cross-validated deviance within one standard error of *λ_opt_*, denoted by *λ*_1_*_se_*. A desirable property of the LASSO is that it performs simultaneous variable selection and parameter estimation through the tuning of the univariate shrinkage parameter.

### 2.2. SCAD

The SCAD algorithm (Fan and Li (2001)), like the LASSO, performs simultaneous variable selection and parameter estimation. Unlike the LASSO, the SCAD penalty is not derived from a known parametric prior distribution, however, the equivalent log-prior can be expressed as

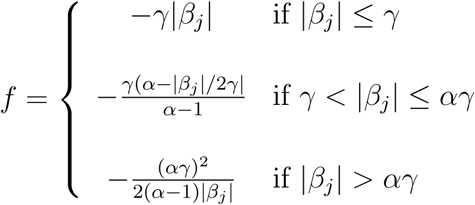

The result of this penalty is to shrink larger regression coefficients less than smaller coefficients relative to the LASSO. The R package ncvreg (Breheny and Huang (2011)) is used for model estimation.

### 2.3. Parallel Regularised Regression Model Search

PReMS fits logistic regression Gaussian shrinkage models to multiple subsets of covariates. The method first fits all possible models with one and two covariates and ranks them based on their log-likelihood. The top *S* two covariate models (user setting with default value of 100) are taken forward to the next stage in which the algorithm determines the unique set of three covariate models that can be constructed by the addition of one covariate to the top two covariate models. The log-likelihood of these models is calculated, and the process continues taking forward the top *S* models to construct models one covariate larger. The user has the option of only searching those two covariate models that can be constructed from the top one covariate models.

The problem can then be split in two: 1) selecting the shrinkage parameter and 2) criteria for selecting the best model.

#### 2.3.1. Selecting the shrinkage parameter

Shrinkage is induced by assigning zero centred Gaussian prior distributions. We parametrise the prior by precision parameter *τ* =1/variance, the log-prior can then be expressed as

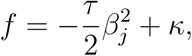

for all coefficients *β_j_*, *κ* is a constant. The prior precision can be interpreted as the penalty parameter in classical penalised regression.

The optimal shrinkage, *τ,* is dependent on the structure of data set under analysis. We tackle this problem by utilising the LASSO estimates *λ_opt_* and *λ*_1_*_se_* and their respective model fits of the data. We then set *τ* by equating the shrinkage induced by the LASSO with the shrinkage induced by the Gaussian prior on the largest regression coefficient of the optimal LASSO fit (*β_max_*)

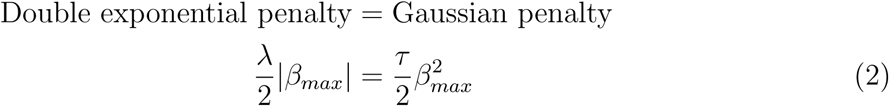

using the optimal LASSO fit and its shrinkage on its largest coefficient *β_max|opt_*

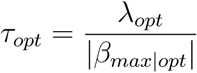

and similarly for the *λ*_1_*_se_* fit to give an estimate

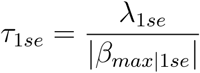

Equating the shrinkage on the largest regression coefficient results in relatively less shrinkage on the other covariate effects. We use the LASSO penalty as it accounts for both the shrinkage of the selected parameters and the number of potential models in the sample space. The appropriate degree of shrinkage can be assessed by examining the calibration slope; this is the slope of a linear regression of the binary outcome against the predicted probabilities, for a well calibrated model this will equal one. With too much shrinkage the predictions are overly shrunk to the mean, the proportion of true positives in the training set, and the slope is greater than one, with too little shrinkage the predictions are overly confident taking values closer to zero and one, resulting in a slope less than one. We show that the PReMS and LASSO have similar calibration slopes for values of *τ* and *λ* satisfying (1).

#### 2.3.2. PReMS: Selecting the best model

We use the Watanabe-Akaike Information Criteria (WAIC) to select the optimal model for a given number of predictors Watanabe (2010). The WAIC is a Bayesian information criteria for estimating the out-of-sample expected error and assumes inference is made from the posterior distribution. WAIC is defined as follows

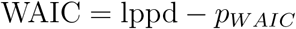

where lppd is the log pointwise predictive density and *p_W AIC_* is a measure of the effective number of parameters included to adjust for over-fitting. lppd and *p_W AIC_* are both defined by expectations over the posterior distribution of the regression coefficients ***β*** as follows

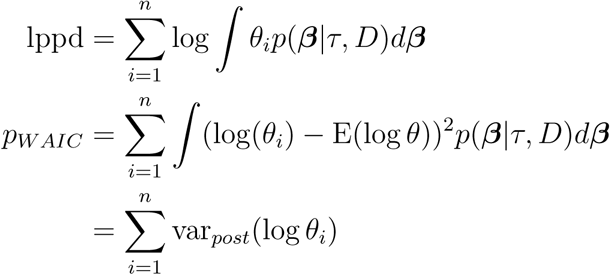

where *D* is the data and *θ_i_* = *p*(*y_i_|****β***). The two quantities are estimated by Monte Carlo integration, taking samples ***β****^s^* from the posterior distribution as follows

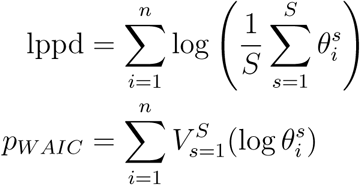

where 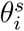 is the evaluation at ***β****^s^* and 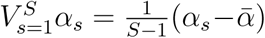. The R package BayesLogit is used to sample from the posterior distribution of the logistic regression coefficients (Polson *et al.* (2013)). Estimation comes at a computation expense and it would infeasible to calculate the WAIC for all models considered. We show in the supplementary material that for a given model size the WAIC is approximately proportional to the log-likelihood, therefore, we only calculate the WAIC for the top ten models ranked by log-likelihood.

We use WAIC rather than the commonly used Akaike information criteria (AIC) since the AIC assumes inference is made at the MLE whereas we make inference from the posterior distribution and thus is suboptimal in settings with strong prior information (Gelman *et al.* (2014)). Furthermore, Gelman *et al.* (2014) show that the WAIC has a more-or-less explicit connection to cross-validation which is computationally expensive but commonly viewed as the gold standard for model selection.

In practice inference is made from the posterior mean of the regression coefficients since, unlike the posterior distribution, the posterior mean allows the model to be simply expressed in terms of fixed regression coefficients, we show for one example data set that this simplification has negligible effect on predictive accuracy. The accompanying R package has a function to make predictions from the posterior distribution.

WAIC is a measure of average prediction error and adjusts for over-fitting by adding a correction for the effective number of parameter, however, it does not account for the number of models and covariates searched over in the PReMS algorithm. Therefore, WAIC is used to determine the optimal model for each model size, but it is not applicable for selecting the optimal model size. Instead cross-validation is used to choose the optimal model size: model fitting is repeated for each training set and applied across all model sizes to the respective test set. PReMS then selects the optimal model size as the one which maximises the mean predictive log-likelihood across all folds. Specifically, if the data is split into *m* folds *J_{_*_1_*_}_,…, J_{m}_*, *k* is chosen to maximise

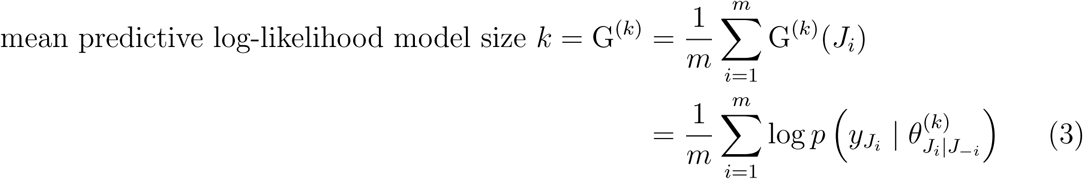

where 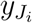 and 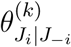 are the outcomes and predictions for each element in the test set *J_i_* given the best fitting model of size *k* determined by training data *J_−i_*. In each fold *τ* is re-estimated given the available training data. Results presented use *m* = 20 folds. We note that in this setting in which many models are searched over WAIC cannot be used to determine the shrinkage parameter *τ* since the effective number of models searched over is dependent on *τ* which is not captured by the WAIC of a single model.

Following this procedure it can occur that many model sizes result in “similar” predictive log-likelihood and it may be somewhat random which of the model sizes is chosen as the “best”. Therefore we also present results for model sizes with predictive log-likelihood within one-standard error of the “best” model – the model with the highest predictive log-likelihood. Following Friedman *et al.* (2010) the standard error of G^(^*^k^*^)^ = (G^(^*^k^*^)^(*J*_1_)*,…,* G^(^*^k^*^)^(*J_m_*)) is calculated as

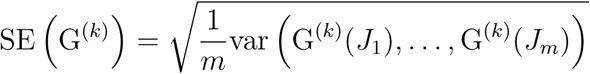

Therefore, in addition to selecting model size *k_max_* (maximising G^(^*^k^*^)^) we select model size *k*_1_*_se_* which is the smallest *k* such that

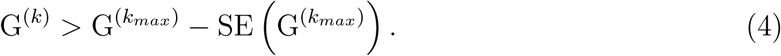

In the text PReMS.opt refers to the model size which maximises G^(^*^k^*^)^ and PReMS.1se refers to the model size sataisfying (4).

#### 2.3.3. Prediction

Throughout, results are presented for inference made at the posterior mean, ie for each sample predict from *θ_i_* = *p*(*y_i_ |* ***β̄***), given in (1), and where

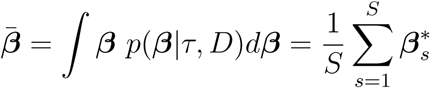

and 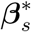 are samples from the posterior. For one data set we also show results for inference made from the posterior mode ***β̂*** and the full posterior distribution of ***β***

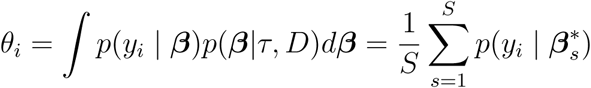

Predictive log-likelihoods calculated from the cross-validation procedure to choose model size (3) were also used to select between *τ_opt_* and *τ*_1_*_se_*.

### 2.4. Data sets

We compare the predictive performance of the PReMS, LASSO and SCAD algorithms applied to six freely available data sets. Of these one is transcriptomic (TB data), two are proteomic (Arcene and Mice), cardiac arrhythmia data and SPECTF Heart data are clinical data sets with alternate measures and the ionosphere data is a physical data set. Description of the data sets are below.

#### 2.4.1. Arcene

The goal of this data is to distinguish ovarian and prostate cancer patients from healthy individuals using protein measures from mass-spectrometry. The available data was split into training and test sets each of 44 cases and 56 controls, with 7000 real probes and 3000 additional noise probes. The data set was prepared by NIPS 2003 as a benchmark data set to compare prediction algorithms (https://archive.ics.uci.edu/ml/datasets/Arcene).

#### 2.4.2. Paediatric Tuberculous

The goal of this data is to distinguish children with tuberculous from children with similar clinical presentations but without tuberculous using 47,323 genome-wide transcriptomic measurements (Anderson *et al.* (2014)). The data is comprised of two cohorts, a discovery set of 135 cases and 89 controls recruited from Malawi and Cape Town and a validation set of 55 controls and 35 cases recruited from Kenya.

#### 2.4.3. Mice Protein Expression data

The goal of this data is to distinguish control mice from trisomic mice (Down syndrome) using 77 measures of protein expression levels. In total there are 570 control samples and 507 case samples (https://archive.ics.uci.edu/ml/datasets/Mice+Protein+Expression).

#### 2.4.4. Cardiac arrhythmia data

The goal of this data is to distinguish cardiac arrhythmia from normal heart function using 191 electrocardiogram measurements (https://archive.ics.uci.edu/ml/datasets/Arrhythmia). In total 245 samples are classified as normal and 206 as abnormal. The abnormal samples are classified into one of 13 abnormal classes but were combined in our analysis.

#### 2.4.5. SPECTF Heart data

The goal of this data is to distinguish between individuals with normal and abnormal heart function using 44 continuous feature pattern. In total 55 individuals are classified as normal and 210 as abnormal (https://archive.ics.uci.edu/ml/datasets/SPECTF+Heart).

#### 2.4.6. Ionosphere data

The goal of this data is to classify radar returns from the ionosphere into good returns and bad from 32 continuous measures (https://archive.ics.uci.edu/ml/datasets/ionosphere). In total 126 returns are classified as bad and 225 as good.

### 2.5. Analyses

All data sets, except Arcene, were randomly split into 100 80% training and 20% test sets, each time ensuring equal proportions of cases and controls in each set. For the TB data the discovery data was randomly split into training and test sets. Measures of model accuracy are calculated in both the test set and Kenyan validation data set for each 100 model fits. In the analysis of this data presented in Anderson *et al.* (2014) only those transcripts with log-fold change *>* 0.5 between cases and controls were taken forward to model building, we follow the same strategy which on average selected 340 transcripts across the training / test splits. A more detailed single analysis of the Arcene data is presented using the available training and test sets. All analyses used 20-fold cross-validation to choose the optimal model size. The following measures were calculated across all test sets:

- number of covariates selected
- area under the receiver operator curve (AUC)
- calibration slope, slope of a linear regression of *y* on *θ*
- root predictive mean square error 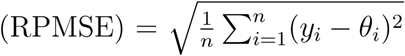

The range of model sizes PReMS searched over was determined by pilot cross-validation analyses such that the predictive log-likelihood was seen to reach a maximum and then decline for increasing model size.

## 3. Results

### 3.1. Analysis of Arcene data

Figure 1 shows both within training set predictive log-likelihood of the Arcene data and test set AUC for increasing model size. For PReMS models fit with both *τ* = *τ*_1_*_se_* and *τ* = *τ_opt_* the relationship between model size and predictive log-likelihood is well mirrored by that between model size and test AUC up to model sizes of 6. For larger models both the predictive log-likelihood and AUC plateau or deteriorate, however, *τ_opt_* displays greater consistency across model sizes. Further Table 1, which displays a summary of accuracy for all methods, PReMS fit with *τ_opt_* has a greater maximum predictive log-likelihood than PReMS fit with *τ*_1_*_se_*.

**Fig. 1.**
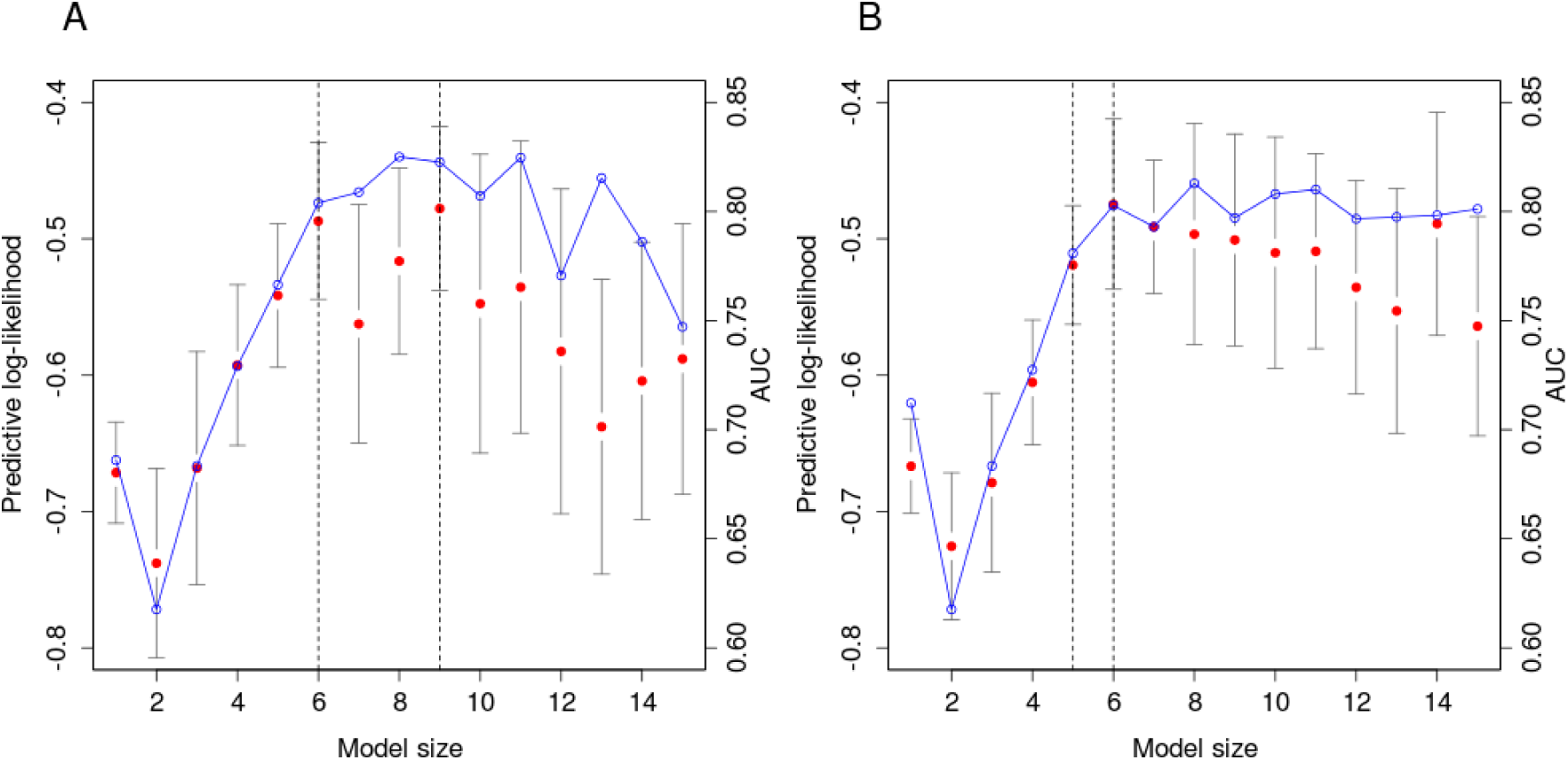
Training set predictive log-likelihood (*•*) and test set AUC (*◦*) by model size for the Arcene data. **A** *τ* = *τ*_1_*_se_*, **B** *τ* = *τ_opt_*. Solid grey bars indicate one standard error of the predictive log-likelihood. Vertical dashed lines indicate the optimal and one standard error fits.

**Table 1:** Results of analyses of the Arcene data by the LASSO, SCAD and PReMS algorithms. *k* is the model size determined by 20-fold cross-validation.

| Method | pll | k | AUC (CI) | RPMSE | Calibration (CI) |
| --- | --- | --- | --- | --- | --- |
| LASSO - 1se | -0.465 | 24 | 0.785 (0.697, 0.873) | 0.433 | 1.11 (0.723, 1.49) |
| LASSO - opt | -0.431 | 31 | 0.783 (0.695, 0.872) | 0.438 | 0.937 (0.598, 1.28) |
| SCAD | -0.518 | 27 | 0.779 (0.690, 0.868) | 0.441 | 0.951 (0.597, 1.30) |
| PREMS- $\tau_{1se}$ - 1se | -0.487 | 6 | 0.804 (0.718, 0.890) | 0.428 | 1.01 (0.679, 1.35) |
| PREMS- $\tau_{1se}$ - opt | -0.478 | 9 | 0.823 (0.742, 0.903) | 0.416 | 1.00 (0.702, 1.30) |
| PREMS- $\tau_{opt}$ - 1se | -0.519 | 5 | 0.781 (0.691, 0.870) | 0.440 | 0.892 (0.568, 1.21) |
| PREMS- $\tau_{opt}$ - opt | -0.475 | 6 | 0.803 (0.716, 0.889) | 0.431 | 0.863 (0.577, 1.15) |

The optimum PReMS model, that which maximises the predictive log-likelihood, occurred with *τ* = *τ_opt_* and six proteins. In comparison, the selected LASSO and SCAD models use between 24 and 31 proteins and have inferior AUC and RPMSE. All models have calibration slope with confidence intervals intersecting 1.

### 3.2. Analyses of other data sets

Table 2 shows the range of model sizes searched over and the proportion of times *τ_opt_* was chosen in favour of *τ*_1_*_se_*, by means of maximising the predictive log-likelihood across the 100 training/test splits. Table 3 summarises the performance of the Methods in terms of average number of markers selected and AUC. For all analyses the optimum PReMS models selected fewer biomarkers than the optimum LASSO models and in the case of the transcriptomic TB test and Kenyan data sets had superior AUC. In all cases PReMS.1se has lower average AUC than PReMS.opt indicating declining performance with smaller model sizes. For the TB test data prems.opt has the highest AUC of all models considered, however, on average prems.opt used one more transcript than Lasso.1se and five more than the SCAD algorithm, although the SCAD method has a greatly reduced AUC. Lasso.1se has the best average AUC in the TB Kenya data, this is likely to reflect heterogeneity between the training and Kenyan data sets and therefore the need for greater shrinkage in training. For comparison, the analysis of the TB data presented in Anderson *et al.* (2014) applied the elastic net to an 80/20 split of the discovery data resulting in a 51 transcript signature with test AUC of 0.862 (95% CI (0.771, 0.940)) and an AUC in the Kenyan of 0.890 (95% CI (0.823, 0.949)).

**Table 2:** *K* – range of models searched over for the PReMS analyses and *P* (*τ_opt_*) – proportion of training runs which selected *τ* = *τ_opt_*.

| Data set | $K$ | $P(\tau_{opt})$ |
| --- | --- | --- |
| TB | 5-25 | 0.19 |
| Mice | 15-40 | 0.06 |
| Cardiac | 5-25 | 0.32 |
| Heart | 1-10 | 0.93 |
| Ionosphere | 1-15 | 0.70 |

**Table 3:** Mean number of covariates selected and AUC for analyses of five test data sets by the LASSO, SCAD and PReMS algorithms. Means were calculated accross 100 random 80/20 training/test splits of the data. Results for the TB test and Kenya data use the same prediction models estimated from each of the 100 training data sets. PReMS.opt results are for model size and *τ* (either *τ*_1_*_se_* or *τ_opt_*) choosen within their respective training sets to maximise the predictive log-likelihood.

| Data set | Lasso.1se |  | Lasso.opt |  | SCAD |  | PREMS.1se |  | PREMS.opt |  |
| --- | --- | --- | --- | --- | --- | --- | --- | --- | --- | --- |
| | $\bar{k}$ | $\overline{AUC}$ | $\bar{k}$ | $\overline{AUC}$ | $\bar{k}$ | $\overline{AUC}$ | $\bar{k}$ | $\overline{AUC}$ | $\bar{k}$ | $\overline{AUC}$ |
| TB test | 19 | 0.886 | 38.1 | 0.889 | 15.8 | 0.874 | 10.2 | 0.882 | 20.3 | 0.892 |
| TB Kenya | - | 0.921 | - | 0.903 | - | 0.91 | - | 0.906 |  | 0.911 |
| Mice | 45.1 | 0.989 | 53.5 | 0.991 | 21.1 | 0.982 | 21.5 | 0.986 | 34 | 0.989 |
| Cardiac | 16.9 | 0.818 | 36.1 | 0.819 | 19.1 | 0.806 | 7.8 | 0.799 | 14.9 | 0.805 |
| Heart | 9.5 | 0.847 | 14.2 | 0.852 | 9.2 | 0.858 | 3.2 | 0.843 | 4.6 | 0.85 |
| Ionosphere | 10 | 0.876 | 17.4 | 0.884 | 9.9 | 0.876 | 5.3 | 0.87 | 9.2 | 0.873 |

For the analyses of the proteomic Mice data only the LASSO.opt models had superior AUC to the PReMS.opt models, however, they on on average used almost 20 more proteins improving the average AUC by 0.002. PReMS.opt and LASSO.1se analyses had the same average AUC, but PReMS.opt used on average 11 fewer proteins. The SCAD analyses had the the lowest average AUC using on average the same number of protein as the PReMS.1se analyses.

For the analyses of the Heart data PReMS analyses frequently identified significantly smaller models than the other methods and on average had only marginally inferior AUC to LASSO.opt models. SCAD models had the highest average AUC but used on average twice as many biomarker compared with PReMS.opt.

PReMS analyses performed worst on the analyses of the Cardiac and Ionosphere data sets. LASSO.opt analyses of the Ionosphere data had significantly better average AUC of all other methods potentially indicating the contribution of many small effects. Models from both LASSO analyses performed well on the Cardiac data potentially indicating that the double exponential prior is most of appropriate of those considered for this data set.

Supplementary Figs 1–6 show boxplots of the number of biomarkers selected, AUC, calibration slope and RPMSE across the 100 training/test splits of the data sets in Table 2. The calibration slopes of the PReMS.*τ_opt_* and PReMS.*τ*_1_*_se_* fits mirror those of Lasso.opt and Lasso.1se respectively indicating the appropriateness of the method for estimating *τ*. In line with LASSO results PReMS.*τ_opt_* had better calibration than PReMS.*τ*_1_*_se_*. There were significant differences in AUC and RPMSE between PReMS.*τ_opt_* and PReMS.*τ*_1_*_se_* across the five data sets with the optimum value varying between data sets, see also Table 2.

### 3.3. Mean, mode and full posterior inference

Table 4 shows a comparison of predictive accuracy using prediction based on the posterior mean, mode and the full posterior distribution across the 100 training/test splits of the mice data. For both the PReMS.*τ*_1_*_se_* and PReMS.*τ_opt_* analyses full posterior prediction resulted in marginally better prediction in terms of the AUC. As would be expected, full posterior prediction, with the encumbant increase in parameter uncertainty, resulted in an increase in the calibration slope. RPMSE captures both factors influencing both the AUC and calibration and shows no clear pattern.

**Table 4:** Comparison of PReMS model fit to the mice data by the posterior mean *β̄*, mode *β̂* and the full posterior *p*(*β|D*).

| | PREMS.$\tau_{1se}$ | | | PREMS.$\tau_{opt}$ | | |
| --- | --- | --- | --- | --- | --- | --- |
| | $\bar{\beta}$ | $\hat{\beta}$ | $p(\beta\vert D)$ | $\bar{\beta}$ | $\hat{\beta}$ | $p(\beta\vert D)$ |
| AUC | 0.9886 | 0.9886 | 0.9892 | 0.9894 | 0.9892 | 0.9899 |
| RPMSE | 0.1737 | 0.1764 | 0.1771 | 0.1776 | 0.1793 | 0.1781 |
| Calibration | 1.026 | 1.038 | 1.044 | 0.989 | 0.998 | 1.004 |

## 4. Discussion

PReMS is more computationally intensive than the SCAD and LASSO algorithms, however, its use of the multiple cores available in desktop computers and the use of computationally efficient methods for selecting the shrinkage parameter *τ* and the optimum model of a given size make the exhaustive model search used in PReMS feasible. However, the cross-validation used to select model size is computationally expensive. Further, choosing model size by maximising the predictive log-likelihood can be quite unstable as shown by the range of model sizes selected in Supplementary Figures 1–6. This occurs as the predictive log-likelihood plateaus and then for increasing model size fluctuations can just be due to random error, see Figure 1. For the application of PReMS to real bio-signature discovery the number of cross-validation folds may have to be increased from the 20 used here to reduce the error in the predictive log-likelihood. Plots of the predictive log-likelihood and its standard error against model size (for example Figure 1) will indicate the stability of the model size estimate. Further there maybe a better method for selecting model size which accounts for both the predictive log-likelihood and its standard error.

The PReMS results presented here could be improved further if a greater range of shrinkage values *τ* was searched over by cross-validation, however, this may be at the expense of model calibration as we have shown that *τ_opt_* typically results in good model calibration.

We have demonstrated good performance of the PReMS method, in comparison with the LASSO and SCAD methods, in the analyses of the the transcriptomic TB data set and the Arcene and Mice proteomic data sets, in terms of both model predictive accuracy and the number of biomarkers selected. Therefore, of the methods compared, PReMS is the most suitable for the discovery of sparse ‘omics biosignatures.

## Footnotes

This preprint was prepared with the AAS LATEX macros v5.0.

## 5. Supplementary Material

Figures S1–6 show comparison of LASSO, SCAD and PReMS model fits applied to 100 random training/test splits for five example data sets. Throughout Lasso.opt is the fit attained from the LASSO penalty parameter that minimises the cross-validated deviance. LASSO.1se is fit attained from the LASSO penalty parameter with cross-validated deviance within 1 standard error of the minimum deviance. PMS.*τ*_1_*_se_*.opt and PMS.*τ*_1_*_se_*.1se are the PReMS fits with penalty parameter derived from the LASSO.1se penalty parameter with the optimal model size and the smallest model size within one standard error of the optimum. Similarly for PMS.*τ_opt_*.opt and PMS.*τ_opt_*.1se.

---

**Fig. S1.**
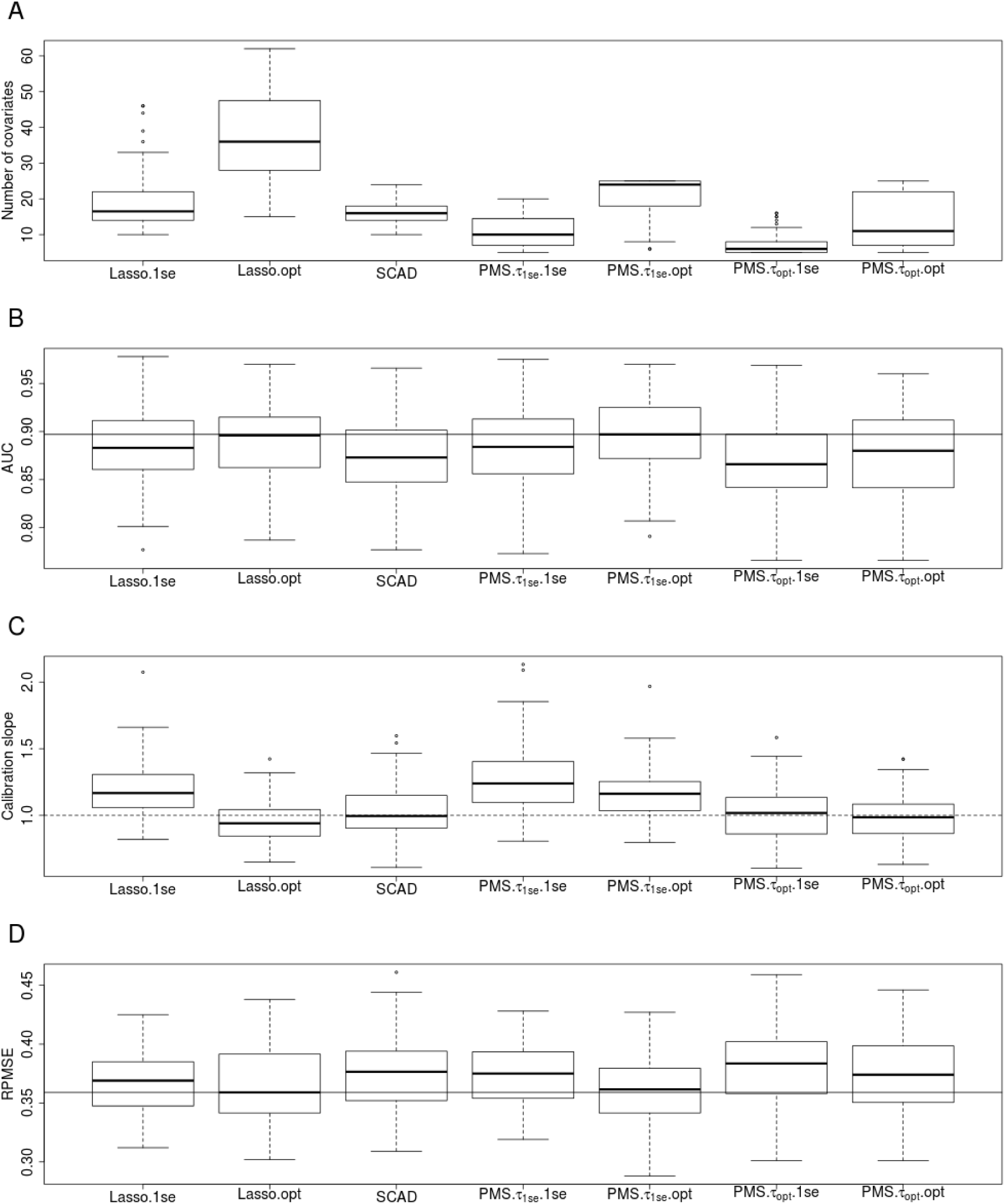
Application to the TB test data set. **A** – number of covariates selected, **B** – area under the receiver operator curve (AUC), **C** – calibration slope, **D** – root predictive mean square error (RPMSE). Dash horizontal lines in plots **B** and **D** indicate the performance of the best performing method. Dash horizontal line in **C** is at one indicting optimum calibration.

**Fig. S2.**
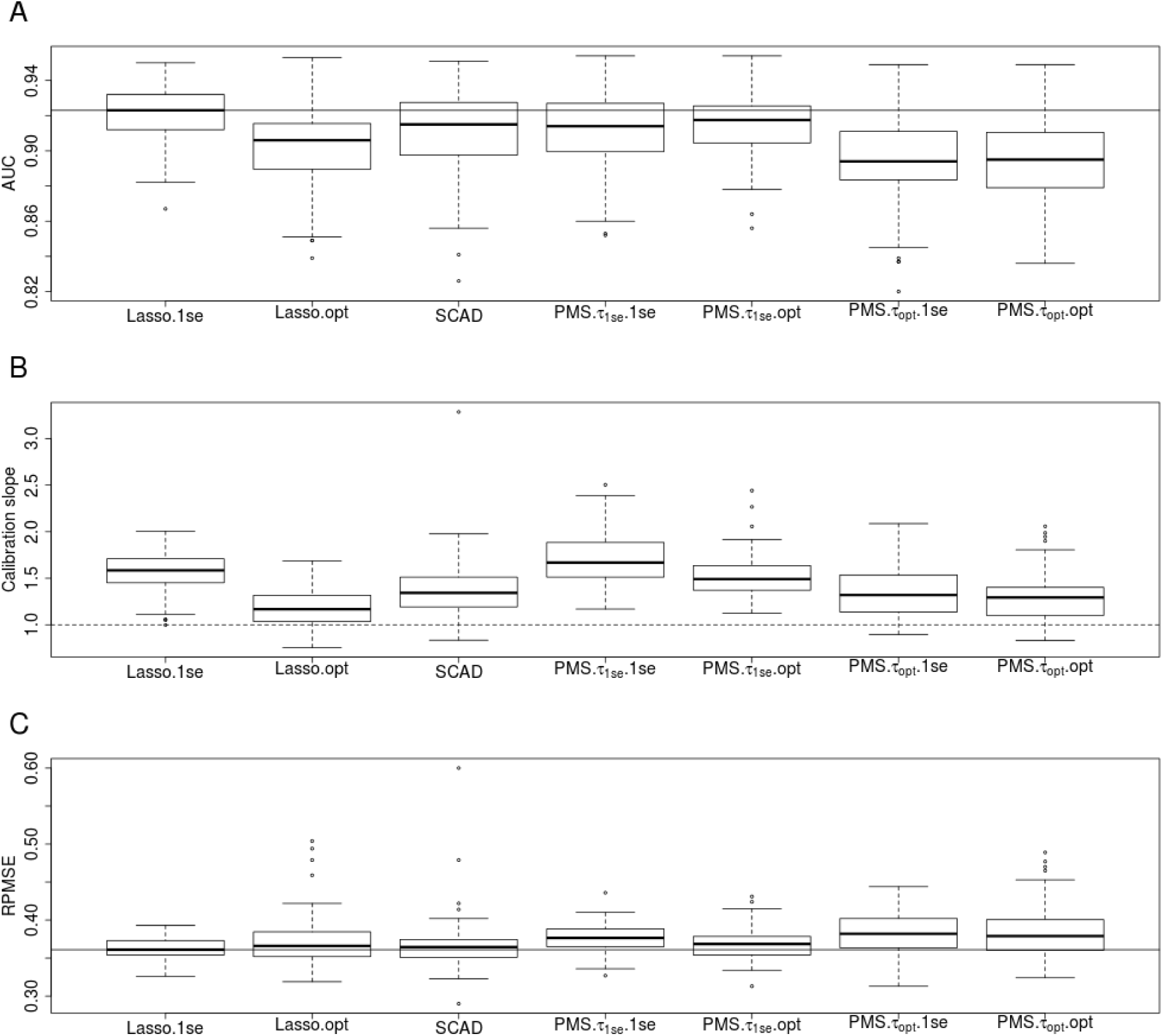
Application to the external Kenya TB validation data. **A** – number of covariates selected, **B** – area under the receiver operator curve (AUC), **C** – calibration slope, **D** – root predictive mean square error (RPMSE). Dash horizontal lines in plots **A** and **C** indicate the performanc eof the best performing method. Dash horizontal line in **B** is at one indicting optimum calibration.

**Fig. S3.**
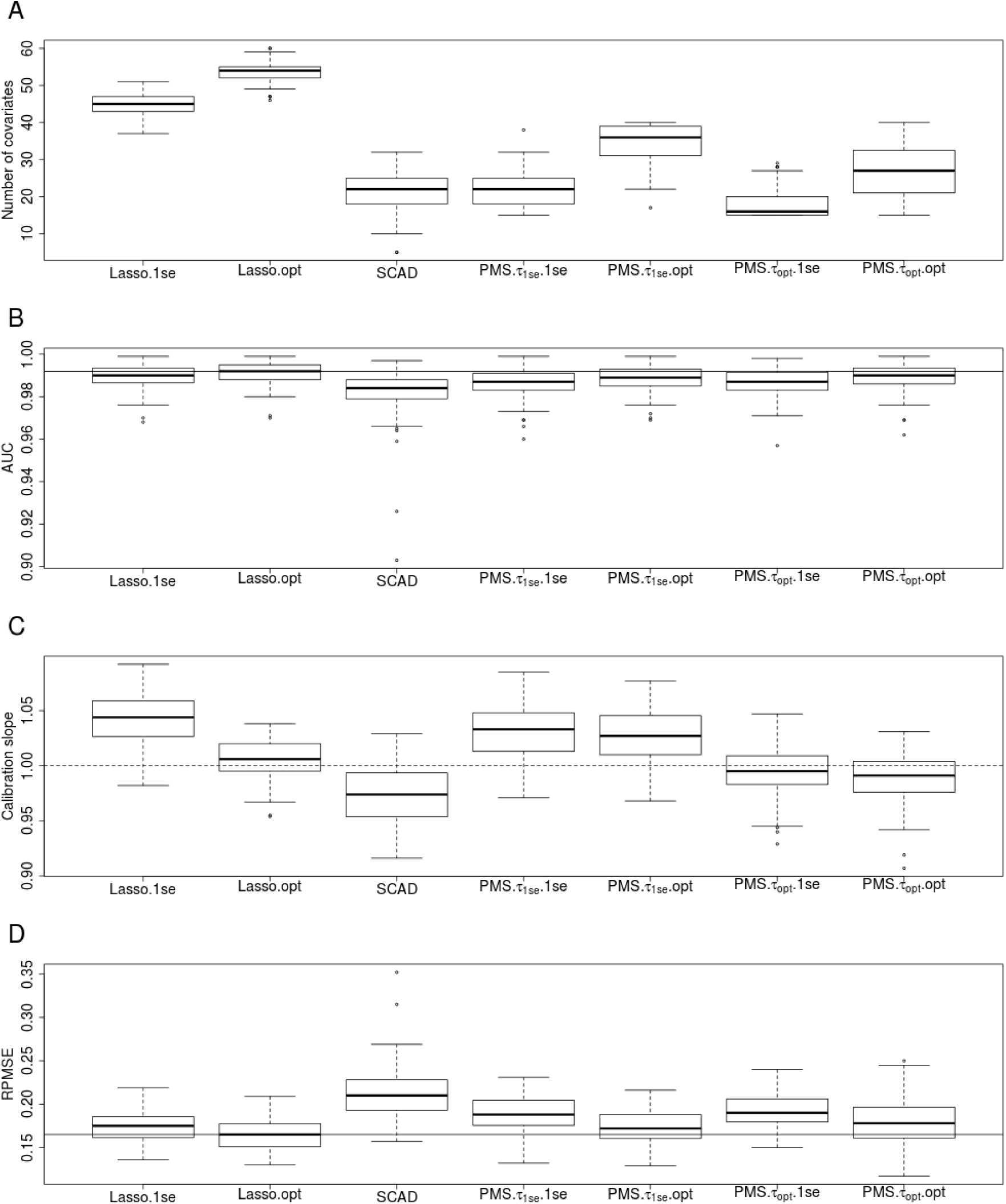
Application to the mice data. **A** – number of covariates selected, **B** – area under the receiver operator curve (AUC), **C** – calibration slope, **D** – root predictive mean square error (RPMSE). Dash horizontal lines in plots **B** and **D** indicate the performance of the best performing method. Dash horizontal line in **C** is at one indicting optimum calibration.

**Fig. S4.**
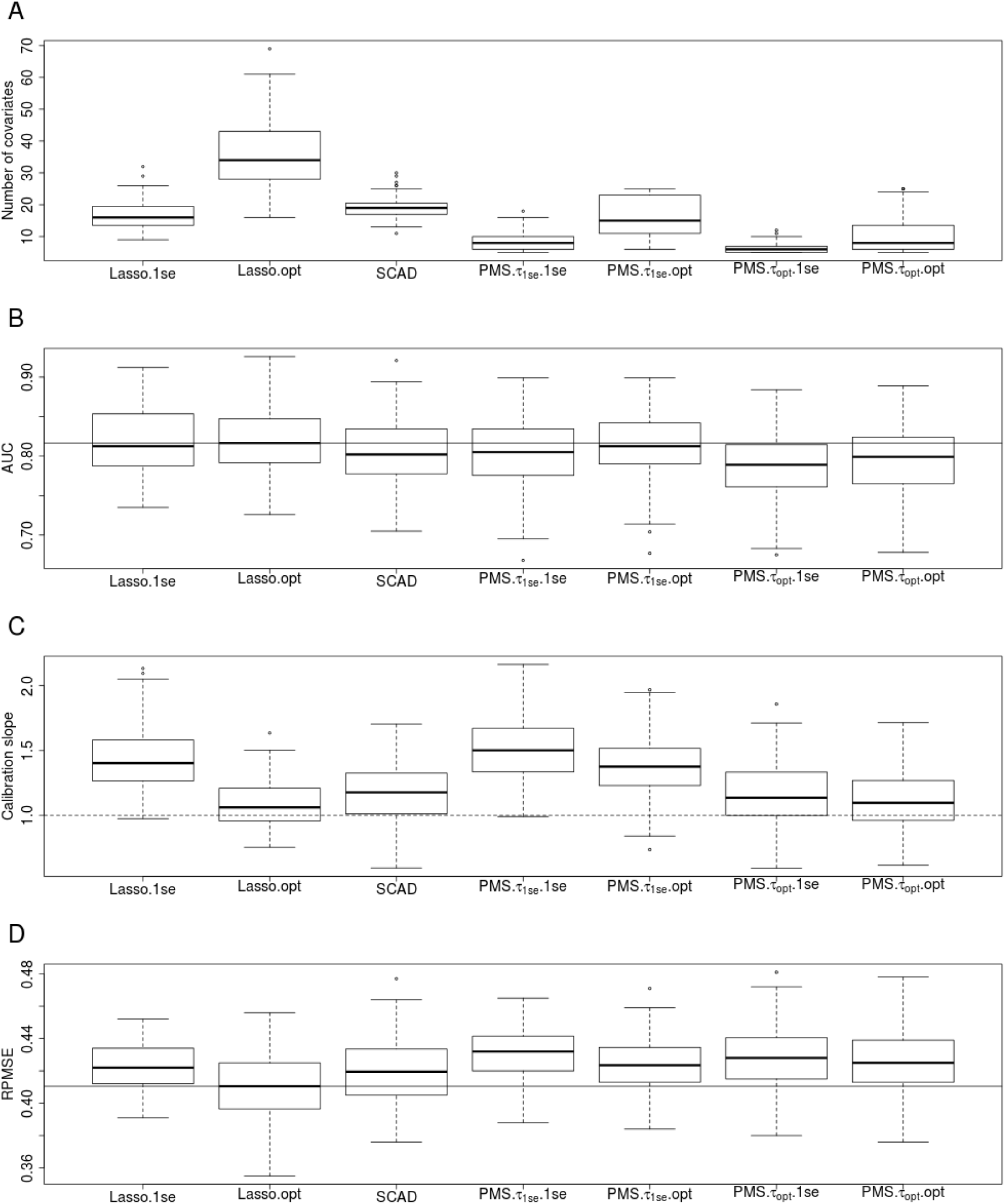
Application to the cardiac test data. **A** – number of covariates selected, **B** – area under the receiver operator curve (AUC), **C** – calibration slope, **D** – root predictive mean square error (RPMSE). Dash horizontal lines in plots **B** and **D** indicate the performance of the best performing method. Dash horizontal line in **C** is at one indicting optimum calibration.

**Fig. S5.**
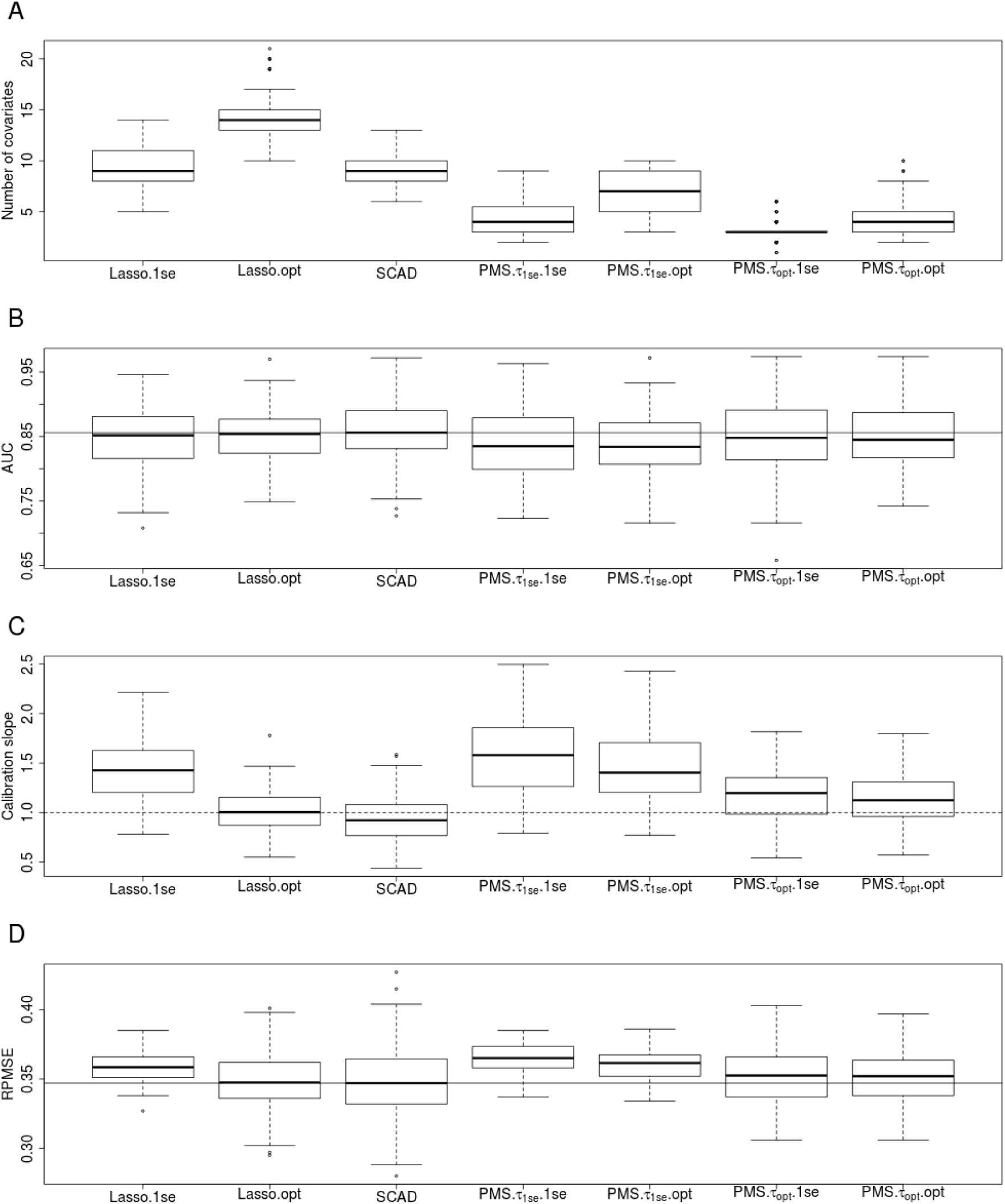
Application to the SPECTF Heart data. **A** – number of covariates selected, **B** – area under the receiver operator curve (AUC), **C** – calibration slope, **D** – root predictive mean square error (RPMSE). Dash horizontal lines in plots **B** and **D** indicate the performance of the best performing method. Dash horizontal line in **C** is at one indicting optimum calibration.

**Fig. S6.**
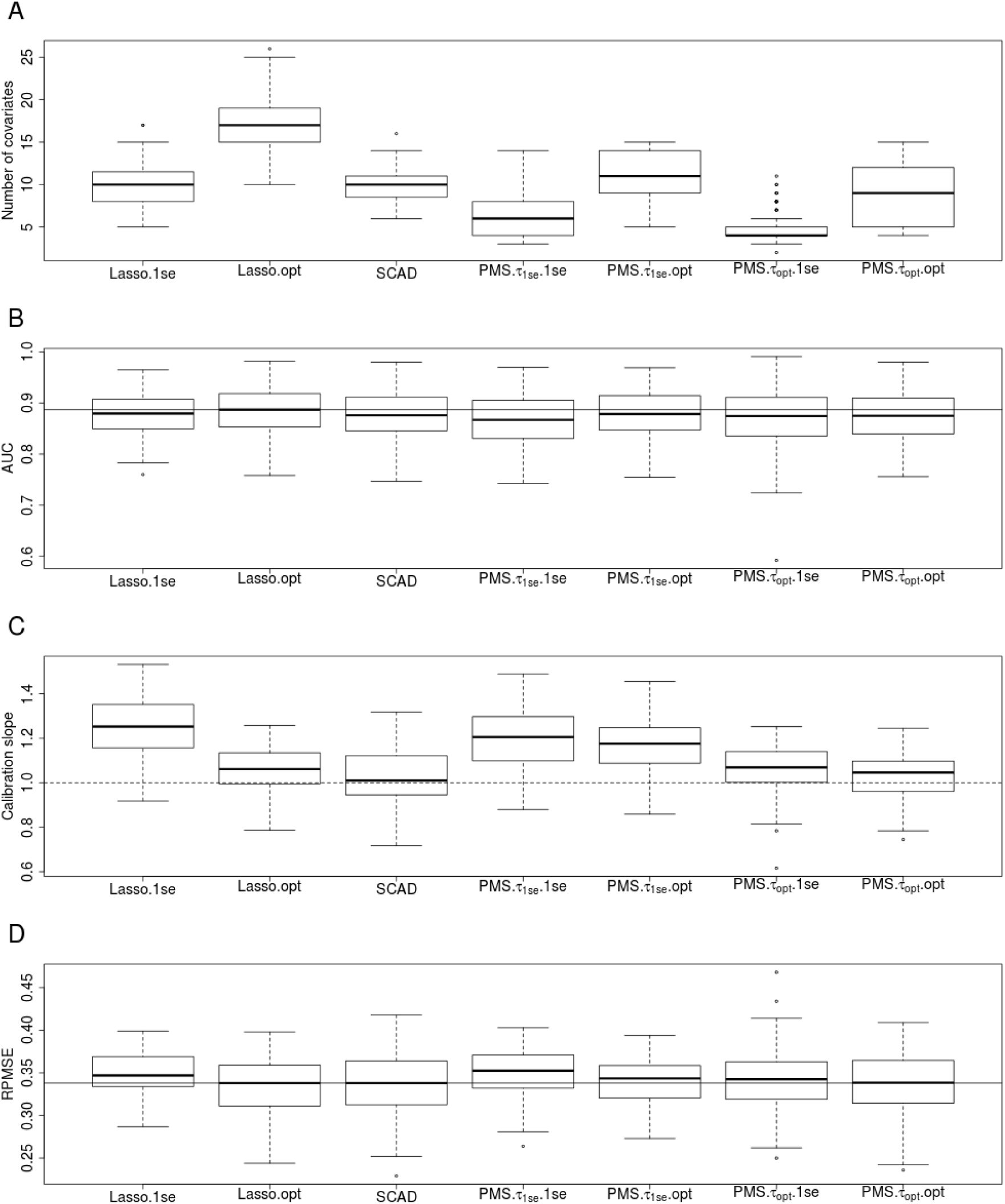
Application to the ionospehere data. **A** – number of covariates selected, **B** – area under the receiver operator curve (AUC), **C** – calibration slope, **D** – root predictive mean square error (RPMSE). Dash horizontal lines in plots **B** and **D** indicate the performance of the best performing method. Dash horizontal line in **C** is at one indicting optimum calibration.

